## Supplementary material for "Merizo: a rapid and accurate domain segmentation method using invariant point attention": Supp. Figure

#### SUPPLEMENTARY DOCUMENT

**Andy M. Lau**
Department of Computer Science
University College London
London, WC1E 6BT
United Kingdom


**Shaun M. Kandathil**
Department of Computer Science
University College London
London, WC1E 6BT
United Kingdom


**David T. Jones**
Department of Computer Science
University College London
London, WC1E 6BT
United Kingdom


May 11, 2023

### 5 Supplementary Methods

#### 5.1 Learning to cluster residues via embedding affinity

The goal of domain segmentation is to assign to each residue $r_i$, a label $k_i$, which allows residues belonging to the same domain to be grouped via label $k$. A property of label $k$ is that it is in a quotient space, that is, the exact value of index $k$ is not important and all labels are equivalent, so long as residues belonging to the same domain end up sharing the same label [1, 2]. Forcing a domain to use a particular label increases the difficulty of the segmentation task, as similar domains may inadvertently have conflicting labels. Several methods have been devised to encourage label-agnostic assignment, one of which is to use an index-invariant learning objective such as via affinity learning [1]. Under this learning regime, and in the context of protein domain segmentation, the embeddings of residues belonging to the same domain are encouraged to be similar to one another when measured by a metric such as cosine similarity, and different when part of different domains [2, 3]. Calculating the pairwise similarity (i.e. affinity) between all pairs of positions, yields an affinity map, $A$, of shape $[N, N]$ (where $N$ is the length of the protein). The ground truth domain arrangement can also be represented by $[N, N]$ domain map $R$ which can be constructed given pairs of residues $r_i$ and $r_j$ as,

$$R_{ij} = \begin{cases} 1, & \text{if } r_i, r_j \text{ belong to the same domain} \\ 0, & \text{otherwise} \end{cases} \tag{1}$$

During training, the goal of the network is to minimise the difference between objective $R$ and the predicted map $A$, by maximising or minimising the similarity of residue embeddings such that $A_{ij} \approx R_{ij}$. Residues with similar embeddings will naturally produce similar probability distributions and by extension, result in the same domain indices being assigned, thus allowing residues to be clustered into domains in a label-agnostic manner.

#### 5.2 Loss functions

The loss of *positive* and *negative* residue pairs (where $R_{ij}$ is equal to 1 and 0, respectively; Equation 1), can be calculated as,

$$\mathcal{L}^{+}_{ij} = 1 - \mathcal{S}_{ij} \tag{2}$$

$$\mathcal{L}^{-}_{ij} = \mathcal{S}_{ij} \tag{3}$$

where $\mathcal{S}_{ij}$ is the cosine similarity between the embeddings of residues $r_i$ and $r_j$. To guide affinity map $A$ towards domain map $R$, we calculate the overall loss of a single input protein using a balanced squared-affinity (BSA) loss, formulated as,

$$\mathcal{L}_{BSA} = \mathcal{L}^{+2} + \mathcal{L}^{-2} \tag{4}$$

The equal contributions of $\mathcal{L}^{+}$ and $\mathcal{L}^{-}$ terms attempt to balance the push and pull forces on the embeddings which leads to faster convergence, while squared terms emphasise the loss when deviations between $A_{ij}$ and $R_{ij}$ are large. The loss of each minibatch is calculated as the sum of four components,

$$\begin{aligned} \mathcal{L}_{total} = \beta_1 \cdot \mathcal{L}_{IPA,BSA} + \beta_2 \cdot \mathcal{L}_{decoder,BSA} \\ + \mathcal{L}_{bg,CE} + \mathcal{L}_{conf,MSE} \end{aligned} \tag{5}$$

where $\mathcal{L}_{IPA,BSA}$ and $\mathcal{L}_{decoder,BSA}$ correspond to the BSA loss applied to the post-IPA single representation of shape $[N, 512]$ and post-decoder domain mask tensor of shape $[N, k]$ and $\beta_1$ and $\beta_2$ are their respective weights (a hyperparameter set to 1 during initial training and 2 during fine-tuning). $\mathcal{L}_{bg,CE}$ is a cross-entropy loss term applied to the non-domain residue predictions of shape $[N, 2]$. $\mathcal{L}_{conf,MSE}$ is the mean squared error (MSE) loss applied to the confidence score predictions and is trained to return values close to 1 when a residue is assigned to the correct domain, and 0 otherwise.

##### 5.3 Evaluation Metrics

###### 5.3.1 Domain pairing

The intersect-over-union (IoU) and Matthew's correlation coefficient (MCC) scores used in our study are first calculated on a per-domain basis and reported as domain length-weighted averages for each target. All scores are determined from the ground truth domain labels, $l_g$, and the predicted domain labels, $l_p$, both of which are vectors of length $N$, where $N$ is the number of residues in a target. Each element in $l_g$ and $l_p$ corresponds to the domain instance label of a residue, and all residues in the same domain will have the same label. The unique set of non-zero labels in $l_g$ and $l_p$ is given by $L_g$ and $L_p$, and $|L_g|$ and $|L_p|$ provide the number of domains in the ground truth and predictions, respectively. As our network is trained in a label-agnostic manner, $L_g$ and $L_p$ will not necessarily match, and the first exercise is to determine the correspondence between them by attempting to assign each value in $L_p$ to one in $L_g$. We obtain this mapping by iterating over the values of $L_g$ (i.e. iterating over each ground truth domain), and obtaining the modal value of $l_p$ predicted for residue positions where $l_g$ is equal to $L_{g,i}$. Each value of $L_g$ can only be assigned to a single value in $L_p$, and when no assignment can be made, the domain is given a score of 0. For example, this can arise when a two-domain protein is predicted as a single domain, making one domain unassignable.

###### 5.3.2 Intersect-over-union

The IoU measures the degree of overlap between a predicted domain segment and its equivalent ground truth, where a score of 1 indicates that the two segments perfectly overlap, and 0 indicates no overlap. The overlap between two segments can be treated as a binary prediction task which allows the equation,

$$IoU = \frac{TP}{TP + FP + FN}, \quad \in [0, 1] \tag{6}$$

where TP (true positives) are the number of overlapping residues between the two segments, FP (false positives) is the number of residues in the predicted domain that are not in the ground truth, and FN (false negatives) are the number of residues in the ground truth that are not in the predicted segment.

###### 5.3.3 Matthew's Correlation Coefficient

The IoU is useful for evaluating how well a true domain is represented by a predicted domain, however, does little to quantify the exact boundary positions. For the latter, we use an MCC score applied directly to the boundaries of each domain (Equation 7). A boundary is considered correctly predicted if it falls within $\pm m$ residues of a true boundary, and we use $m$ at values of 5, 10 and 20 residues. For all paired ground truth and predicted domains, we take the expected and predicted boundary indices and use a linear sum assignment algorithm to obtain an optimal 1:1 pairing between the two, while respecting $\pm m$. The confusion matrix for each domain can be defined as:

- True positives - the number of paired ground truth boundaries
- False positives - is the number of unpaired predicted boundaries
- False negatives - is the number of unpaired ground truth boundaries
- True negatives - are all other residue positions

$$MCC = \frac{TP \times TN - FP \times FN}{\sqrt{(TP + FP)(TP + FN)(TN + FP)(TN + FN)}}, \quad \in [-1, 1] \tag{7}$$

###### 5.3.4 Domain count

The deviation between the expected $|L_g|$ and predicted number of domains $|L_p|$ in a single target can be quantified by calculating the absolute error between them. Averaging across a set of targets $\{X\}$ provides the mean absolute error (MAE) metric which summarises the average deviation of the predicted domain count against the ground truth and is given by,

$$MAE = \frac{1}{|\{X\}|} \sum_{i=1} ||L_{p,i}| - |L_{g,i}||, \quad \in [0, \infty] \tag{8}$$

**Supplementary Table 1:** Training and test set statistics.

| | **Training** | **% of total** | **Testing** | **% of total** | **Total** |
|---|---|---|---|---|---|
| **No. Components** | 537 | 82.0 | 118 | 18.0 | 655 |
| of which contain a single superfamily | 62 | 83.8 | 12 | 16.2 | 74 |
| of which contain >1 superfamily | 475 | 81.8 | 106 | 18.2 | 581 |
| **No. Topologies** | 974 | 86.0 | 158 | 14.0 | 1132 |
| **No. Superfamilies** | 3587 | 92.4 | 293 | 7.6 | 3880 |
| **No. Domains** | 43214 | 96.0 | 1780 | 4.0 | 44994 |
| **No. Chains** | 18231 | 96.2 | 725 | 3.8 | 18956 |
| Redundant chains (seqid > 0.99) | 944 | 93.8 | 62 | 6.2 | 1006 |
| **Final No. Chains** | 17287 | 96.3 | 663 | 3.7 | 17950 |

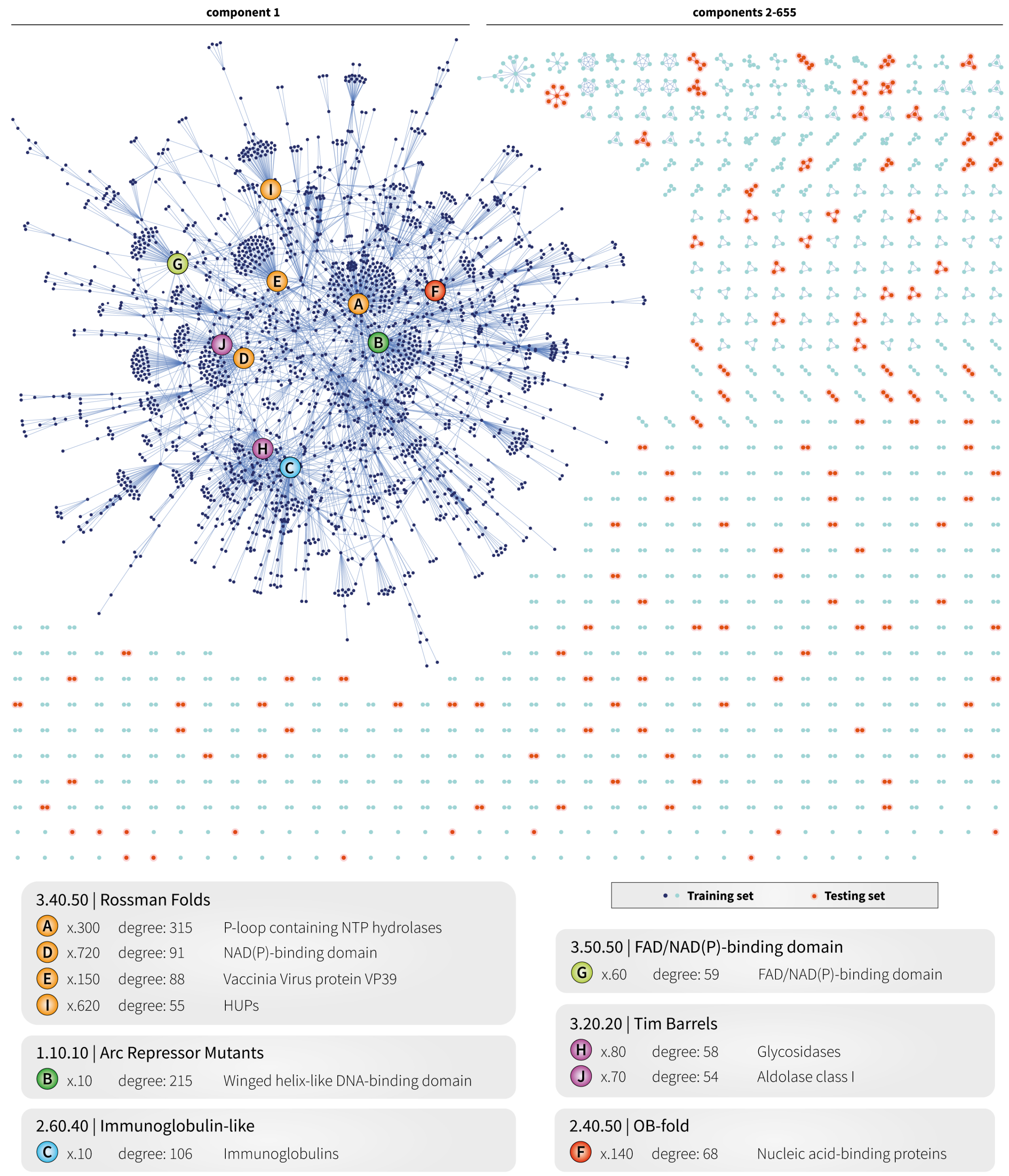


**Supplementary Figure 1: CATH superfamily graph.** The graph is composed of 3880 nodes across 655 components (i.e. sub-graphs) where each node represents a CATH superfamily. Two nodes are connected when a PDB chain containing domains from both superfamilies can be found. The 10 most connected superfamilies are highlighted by labels A-J. A summary of each hub superfamily is shown in the legend at the bottom of the figure. Blue and red components represent those assigned to either the training or test set. The lack of components sharing both green and red nodes indicates that both sets do not overlap at the superfamily level. Additional training and testing set statistics are reported in Supp. Table 1.

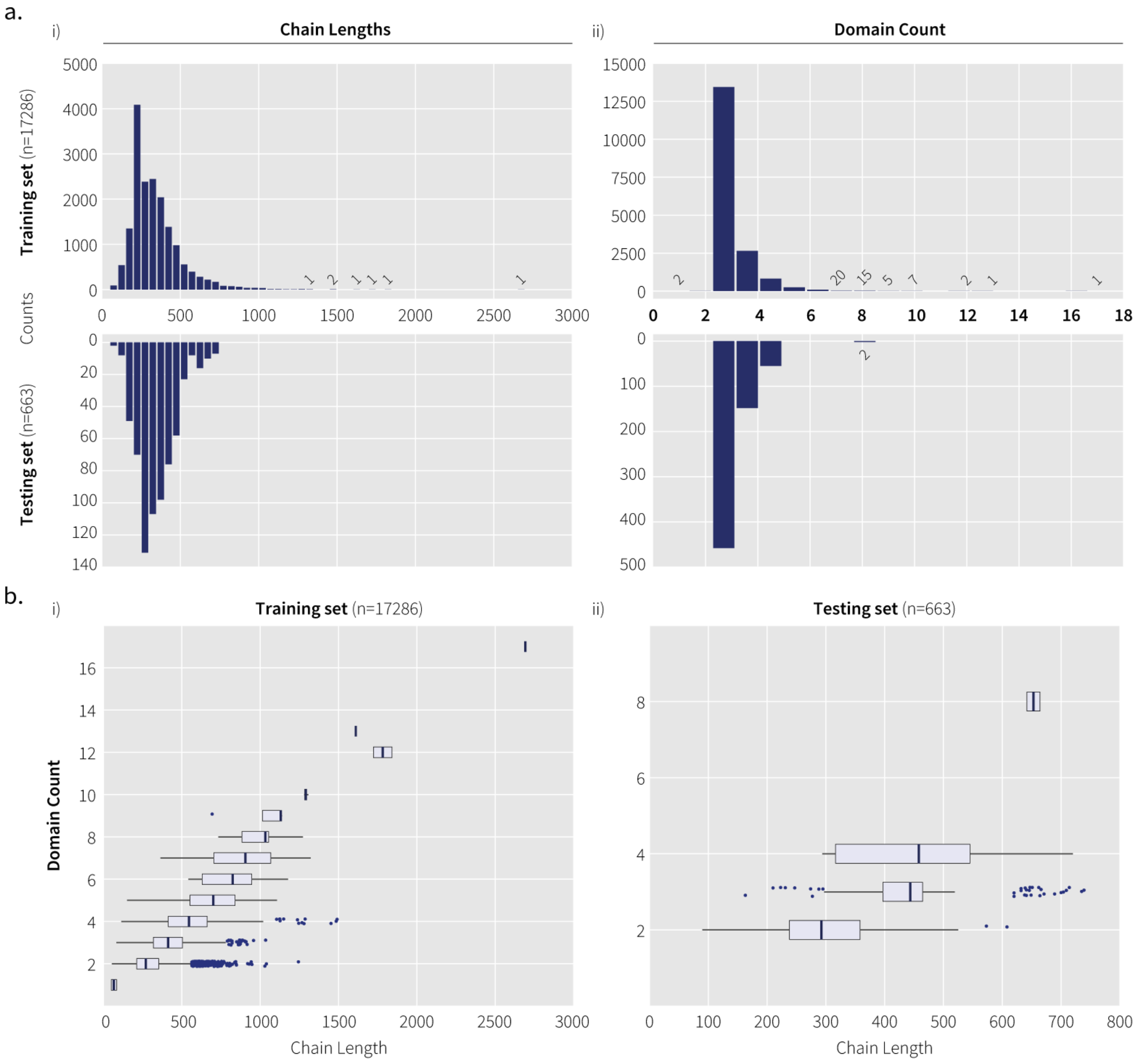


**Supplementary Figure 2: Training and test set statistics.** **a**) Chain lengths and domain count distributions for targets in the training and test sets. Bins with very few members are labelled with the bin count for clarity. **b**) Chain length distributions delimited by domain count for i) training and ii) test sets. The bar within each box represents the distribution median. Outliers are defined as chains with a length exceeding 1.5xIQR. The two targets in the training set with a domain count of 1 result from holding pen domains being removed from multi-domain chains.

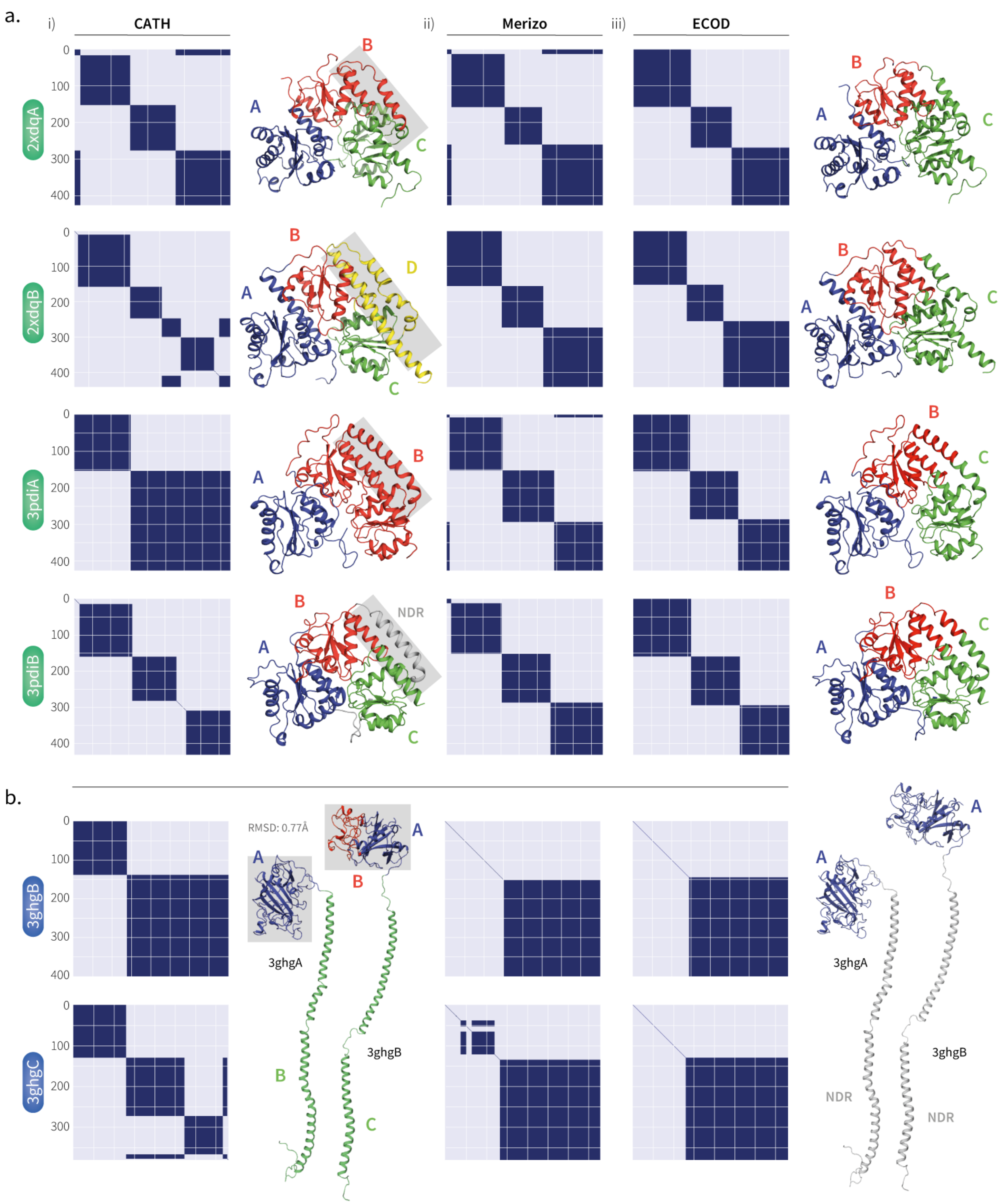


**Supplementary Figure 3: Examples of targets where Merizo agrees with ECOD but not CATH.** The i) CATH domain map and structure, ii) Merizo-predicted domain map and iii) ECOD domain map and structure are shown for two sets of multidomain targets from the CATH-663 set. Set (a) contains the subunit structures of Protochlorophyllide Oxidoreductase (2xdqA and 2xdqB) and precursor-bound NifEN (3pdiA and 3pdiB). All four chains are classified as three domains by ECOD and Merizo but vary in domain assignment by CATH despite having similar structures. The grey box highlights a region of three helices which differ in assignment in each chain. In example (b), the assignment of the C-terminal domain in human fibrinogen (3ghgA and 3ghgB) differs in CATH despite having only a 0.77Å RMSD difference (only residues highlighted by the grey box are aligned). Domain assignments by Merizo and ECOD are mostly identical with the exception of Merizo classifying a portion of the N-terminal tail as a separate domain.

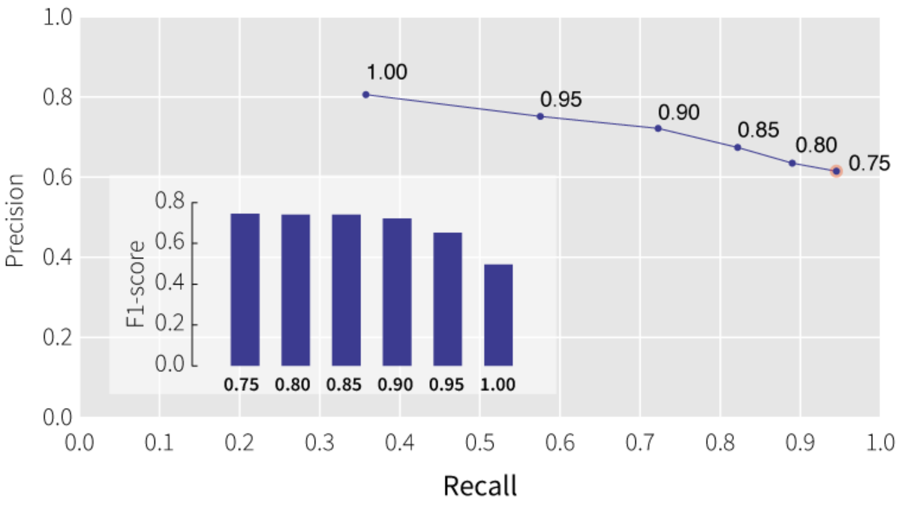


**Supplementary Figure 4: Determining an optimal pIoU threshold** Precision-recall curve showing the precision, recall and F1-score obtained when using the pIoU as a binary measure of domain assignment quality. The highest F1-score obtained is from using a pIoU cutoff of 0.75. Domains with a pIoU of 0.75 or greater are likely true positives, while those below this value are likely false positives.

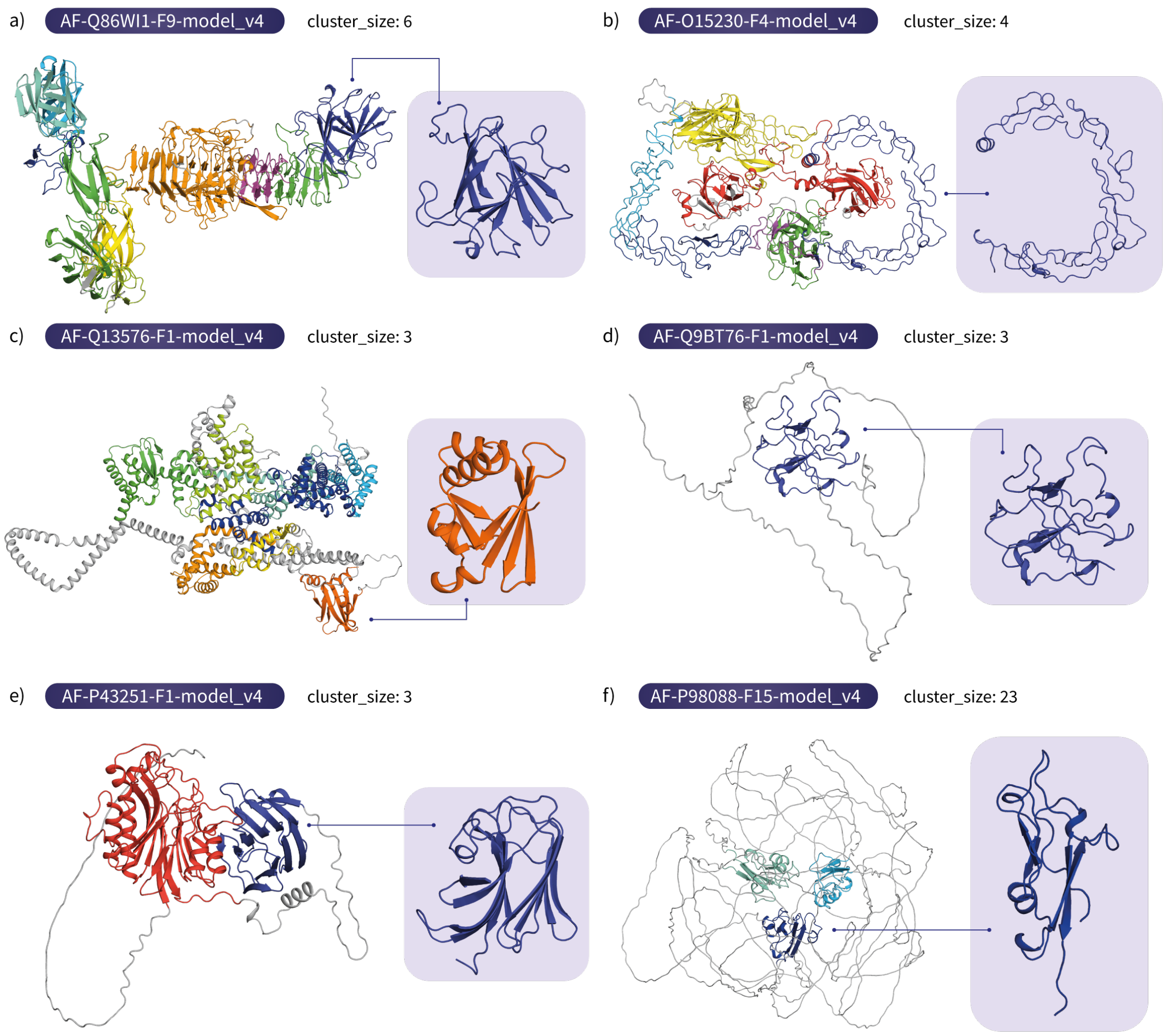


**Supplementary Figure 5: Examples of folds identified in the AFDB-human set not matched to CATH representative domains.** In each panel a-f, the full AFDB model as well as the non-CATH-matched domain is shown. Models shown are the cluster representatives of the subset of domains identified in the AFDB-human set that cannot be aligned to a CATH S40 representative domain based on the SSAP score.

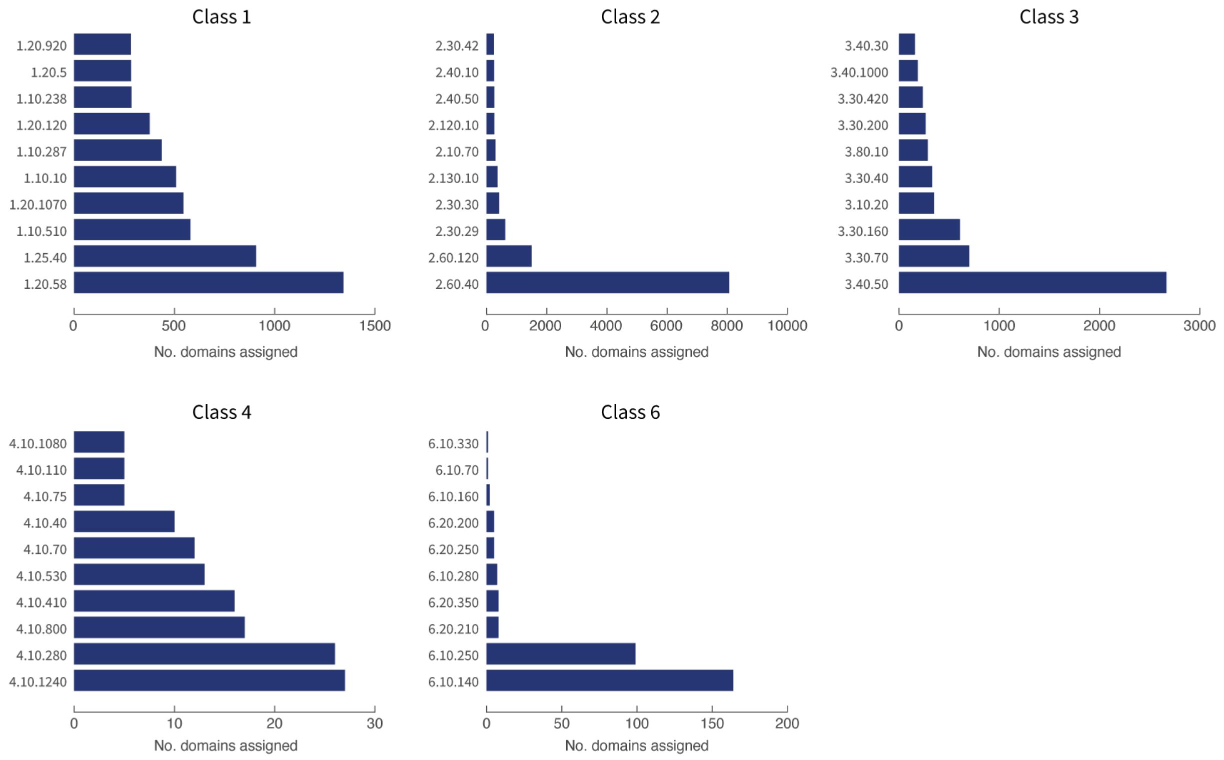


**Supplementary Figure 6: Number of domains putatively assigned to CATH topologies in the AFDB-human set.** The 10 most abundant CATH topologies identified in each CATH class are shown. Data encompasses 34,564 domains that could be matched to CATH S40 representative domains with a SSAP score of at least 70, indicating topology-level similarity.

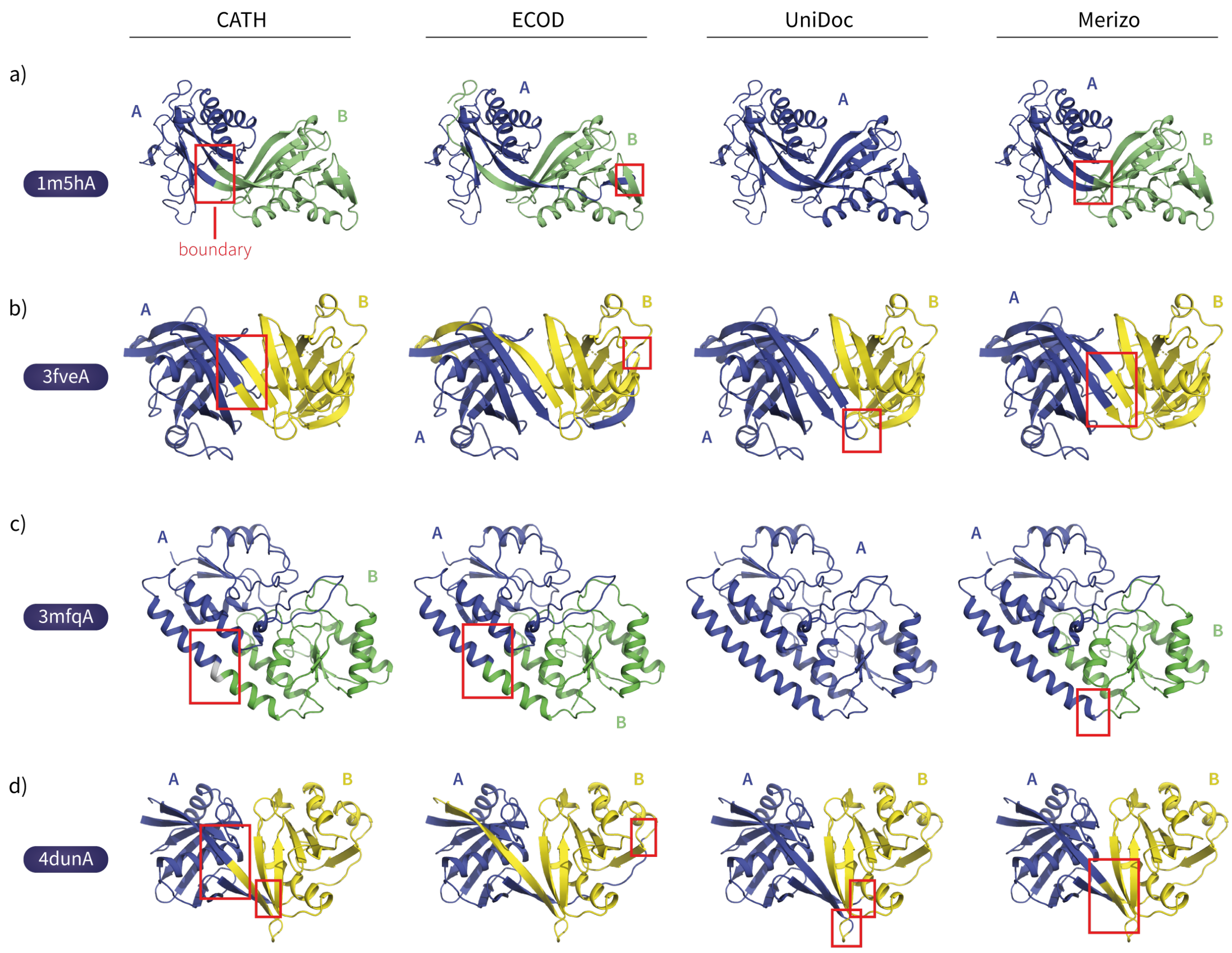


**Supplementary Figure 7: Examples of multi-domain proteins with domain boundaries on secondary structure elements.** Four examples are shown where in each case, the CATH domain boundary assignment is within a secondary structure element (red box). In c) both CATH and ECOD assign a boundary to the middle of a long helix which runs adjacent to the two domains. Domain boundary assignments by UniDoc are restricted to residues not part of secondary structure elements, while Merizo does not have this restriction. In examples b and d, UniDoc assigns the boundary to the end of the nearest $\beta$-strand, while the assignment by Merizo is more precise and is made to the strands themselves.

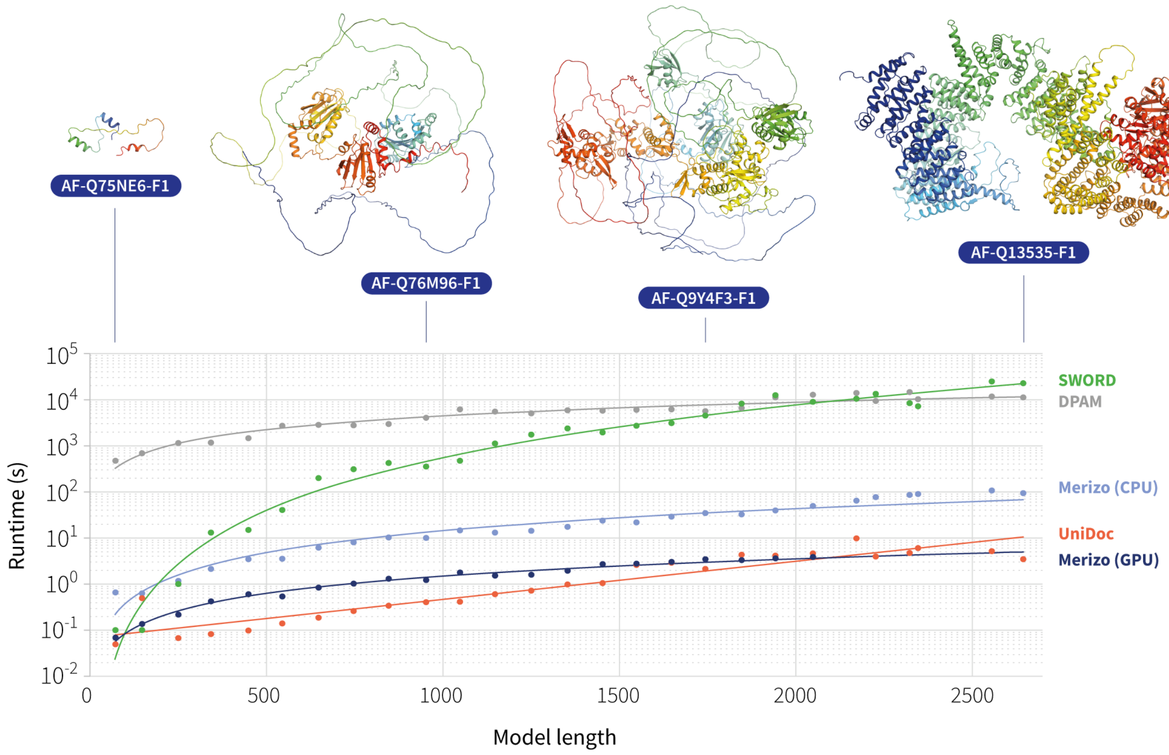


**Supplementary Figure 8: Comparison of runtimes for the AFDB-27 set** The model lengths of the AFDB-human set were divided into 100-residue bins (27 in total) and one model was selected from each bin at random, to form a set of 27 models encompassing the full range of AFDB model lengths. Each method was timed on how long it took to segment models. Merizo timings were conducted on either a single GPU (NVIDIA GTX 1080Ti with 11GB of memory) or a single CPU (Intel Xeon E5-1620 v3 @ 3.50GHz). Line functions of the form $y = bx^k$ were fit to each set of data points with the exception of UniDoc data which was fit to an exponential function. Note that 6 of the longest models were not processed by Merizo (GPU) due to memory limitations.

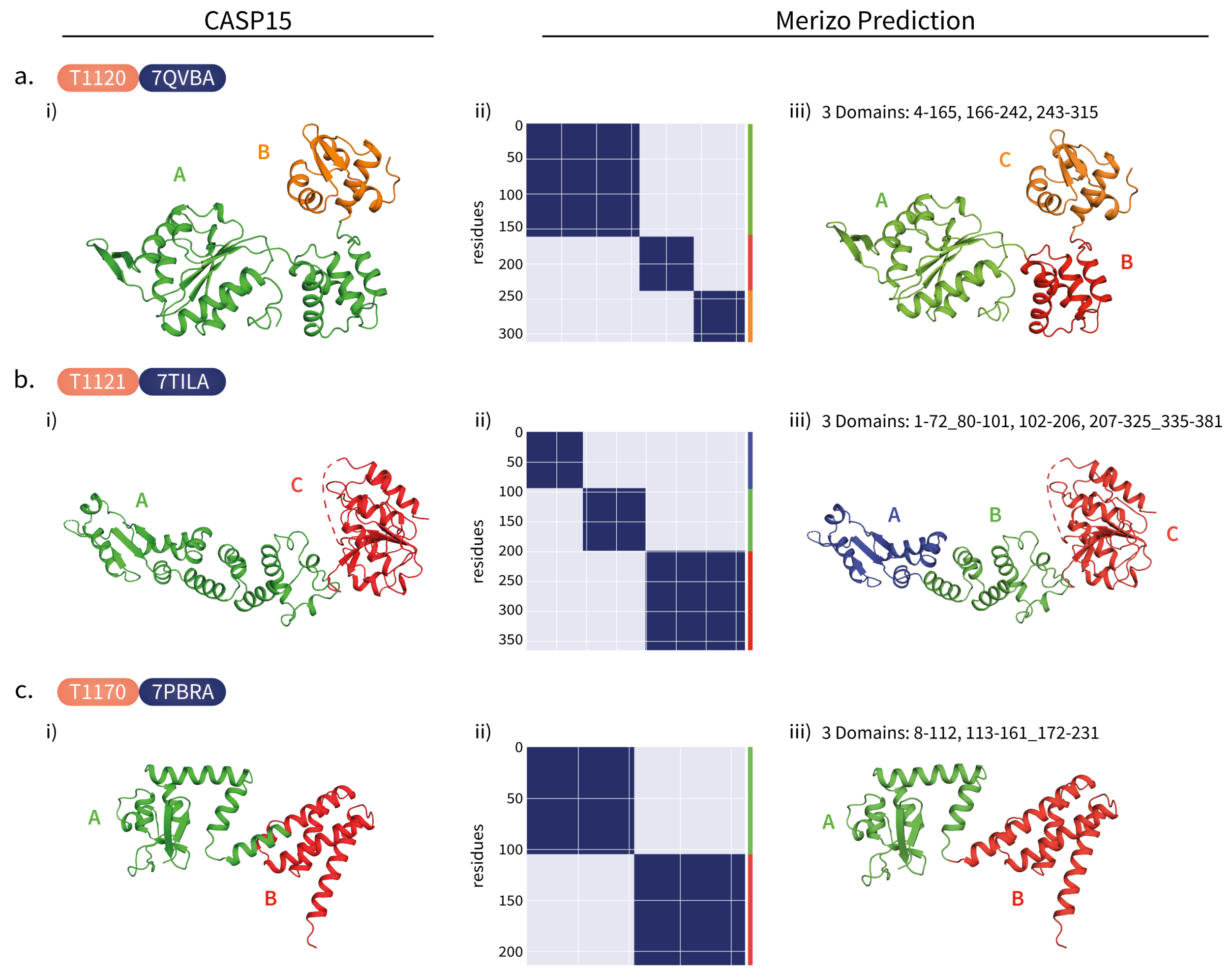


**Supplementary Figure 9: Segmentation maps for CASP15 multi-domain targets.** i) Domain classifications as provided by CASP organisers (left) vs ii) Merizo predictions, are shown for three multidomain targets T1120, T1121 and T1170. Targets T1121 and T1170 are annotated as two-domain proteins in CASP15 but are predicted as three domains by Merizo.

#### References

[1] Long Jin, Zeyu Chen, and Zhuowen Tu. Object detection free instance segmentation with labeling transformations. *CoRR*, abs/1611.08991, 2016.

[2] Yen-Chang Hsu, Zheng Xu, Zsolt Kira, and Jiawei Huang. Learning to cluster for proposal-free instance segmentation. *CoRR*, abs/1803.06459, 2018.

[3] Wei Huang, Shiyu Deng, Chang Chen, Xueyang Fu, and Zhiwei Xiong. Learning to model pixel-embedded affinity for homogeneous instance segmentation. In *Proceedings of the AAAI Conference on Artificial Intelligence*, volume 36, pages 1007–1015, 2022.
